# Increased low-frequency brain responses to music after psilocybin therapy for depression

**DOI:** 10.1101/2022.02.13.480302

**Authors:** Matthew B. Wall, Cynthia Lam, Natalie Ertl, Mendel Kaelen, Leor Roseman, David J. Nutt, Robin L. Carhart-Harris

## Abstract

Psychedelic-assisted psychotherapy with psilocybin is an emerging therapy with great promise for depression, and modern psychedelic therapy (PT) methods incorporate music as a key element. Music is an effective emotional/hedonic stimulus that could also be useful in assessing changes in emotional responsiveness following psychedelic therapy. Brain responses to music were assessed before and after PT using functional Magnetic Resonance Imaging (fMRI) and ALFF (Amplitude of Low Frequency Fluctuations) analysis methods. Nineteen patients with treatment-resistant depression underwent two treatment sessions involving administration of psilocybin, with MRI data acquired one week prior and the day after completion of the second of two psilocybin dosing sessions. Comparison of music-listening and resting-state scans revealed significantly greater ALFF in bilateral superior temporal cortex for the post-treatment music scan, and in the right ventral occipital lobe for the post-treatment resting-state scan. ROI analyses of these clusters revealed a significant effect of treatment in the superior temporal lobe for the music scan only. Somewhat consistently, voxelwise comparison of treatment effects showed relative increases for the music scan in the bilateral superior temporal lobes and supramarginal gyrus, and relative decreases in the medial frontal lobes for the resting-state scan. ALFF in these music-related clusters was significantly correlated with intensity of subjective effects felt during the dosing sessions. These data suggest a specific effect of PT on the brain’s response to a hedonic stimulus (music), implying an elevated responsiveness to music after psilocybin therapy that was related to subjective drug effects felt during dosing.

## Introduction

The use of psychotropic compounds for medicinal, spiritual, and recreational purposes has ancient origins in a diverse set of human cultures, and likely stretches back into pre-history (Hardy, 2021). A recent revival of interest in the clinical potential of these compounds has found that classic psychedelics such as psilocybin may have utility in the treatment of depression (Carhart-Harris et al., 2021; Carhart-Harris et al., 2016), addiction (Johnson, Garcia-Romeu, Cosimano, & Griffiths, 2014), and anxiety (Grob et al., 2011). Modern neuroscientific research is beginning to understand the potential physiological and psychopharmacological mechanisms behind these therapeutic effects (Carhart-Harris et al., 2017; Mertens et al., 2020).

Music is also pervasively, and perhaps universally, enjoyed across cultures and throughout human history (Cross, 2001), and has deep biological and evolutionary foundations (Fitch, 2006). The use of music in a therapeutic context dates back to at least Pythagoras (Nilsson, 2008) and also features in the ancient Indian medical system of Ayurveda (Sundar, 2007). Modern research has started to identify the neurological underpinnings of the unique role that music plays in human cultural and emotional life (Reybrouck, Vuust, & Brattico, 2018) and its potential clinical use (Bower, Magee, Catroppa, & Baker, 2021; Nilsson, 2008). The brain’s reward system, which includes the ventral striatum and ventro-medial pre-frontal cortex (vmPFC), appears to be implicated in pleasurable emotional responses to music (Koelsch, 2020). The primary auditory cortex (superior temporal gyrus, Heschl’s gyrus) is another relevant system for music perception (Koelsch, Skouras, & Lohmann, 2018) and limbic brain regions have also been implicated in emotional responses to music (Koelsch & Skouras, 2014). Music can evoke a range of emotions but generally can be considered a hedonic stimulus, capable of evoking positively valanced emotions. It is therefore a potentially useful tool for indexing anhedonia, i.e. the relative inability to experience pleasure; a common symptom of depression (Cao et al., 2019). Previous studies have shown decreased responses to music in the neural reward system in depressed patients (Jenkins et al., 2018; Osuch et al., 2009).

Recent research on the effect of psychotropic drugs on music perception has shown that classic psychedelics, like LSD, can enhance the subjective emotional response to music (Kaelen et al., 2015), and that brain responses to music are also significantly increased (Kaelen et al., 2017). The hippocampus has been identified as a key region involved in both the effects of LSD and music, alone, and in combination (Kaelen et al., 2016). Music plays a central role in modern and historic versions of psychedelic therapy, useful both for its calming effects at various stages of the therapeutic process, and for its ability to act synergistically with the drug to guide and potentially enhance emotional experiences and evoke autobiographical memories (Barrett, Preller, & Kaelen, 2018; Kaelen et al., 2016).

The aim of the present study was to examine the effects of psychedelic therapy for depression on the brain’s response to music using functional Magnetic Resonance Imaging (fMRI), comparing a music-listening scan to a resting-state scan, before and after treatment. To do so, we used a measure of the brain’s Low-Frequency Oscillations (LFOs) known as ALFF (Amplitude of Low-Frequency Fluctuations; Zang et al., 2007). ALFF is a spatially unconstrained analysis method that is well-suited for characterizing both resting-state and stimulus-related brain activity and has high test-retest reliability (Li, Kadivar, Pluta, Dunlop, & Wang, 2012). We also investigated the relationship between the derived ALFF results and subjective clinical/psychometric results obtained during and after the therapy sessions.

## Methods

This study was approved by the National Research Ethics Service (NRES) committee (West London) and was conducted in accordance with the revised declaration of Helsinki (2000), Good Clinical Practice (GCP) guidelines, and the National Health Service (NHS) Research Governance Framework. Imperial College London sponsored the research which was conducted under a Home Office license for research with schedule 1 drugs, and the Medicines and Healthcare products Regulatory Agency (MHRA) also approved the study. All patients gave written informed consent. The study used facilities at the Imperial College Clinical Research Facility, and Invicro London.

The resting-state data used in this work has been previously analysed and reported (Carhart-Harris et al., 2017) but not using an ALFF analysis. The presently reported results therefore derive from a novel analysis of the resting-state data, combined with the music-listening data from the same subjects. The music-listening fMRI data has not been previously reported.

### Participants and Recruitment

Nineteen subjects were recruited and completed the study, including thirteen males and six females, aged between 27-64. The mean age of participants was 41.3 (SD=10.5), and all had diagnoses of treatment resistant major depression. Initial screening included physical health assessments (electrocardiogram, blood and urine tests), a psychiatric interview, an assessment by a qualified clinician, and self-report questionnaires. The key inclusion criteria were a diagnosis of moderate to severe depression, with a score of 17 or higher on the 21-item Hamilton Depression Rating Scale (HAM-D), and treatment-resistance, meaning that they had been non-responsive to at least two previous pharmacological treatments. Exclusion criteria included previous or current psychotic disorders, individuals with a history of psychotic disorders in their immediate family members, previous suicide attempts that led to hospitalization, pregnancy, drug or alcohol dependence, phobia of blood or needles, history of mania, other concurrent medications, and general contraindications for MRI scanning.

### Design and Procedure

For full details of the study procedure please see the original report of the clinical data (Carhart-Harris et al., 2016). The study was an open-label design with no control group or placebo, and all subjects received the active intervention with full prior disclosure. The psilocybin used in the study was obtained from THC-pharm (Frankfurt, Germany), and processed into size 0 capsules with 5 mg psilocybin each, by Guy’s and St Thomas’ Hospitals’ Pharmacy Manufacturing Unit (London, UK).

Psilocybin was administered to the participants in two therapy sessions: low dose (10mg) in the first session and high dose (25mg) in the second session one week later. Post capsule ingestion, patients lay with eyes closed while listening to music. Two therapists were always present, and adopted a non-directive, supportive approach for the duration of the sessions.

MRI scanning visits were conducted one week prior to the first therapy/dosing visit, and the day after the second therapy/dosing session. The primary clinical outcome was the Quick Inventory of Depressive Symptoms (Rush et al., 2003) and this was administered at baseline, weekly from week one to week five, and finally at a three-month follow-up. Other questionnaire measures included the 5D-ASC (5-dimension altered states of consciousness questionnaire; administered 6-7 hours post-dosing, at each dosing session; Dittrich, 1998) and the GEMS-3 (Geneva Emotional Music Scale; Zentner, Grandjean, & Scherer, 2008), which was administered immediately after each MRI scanning session in order to assess the subjective response to the music-listening scan. For full details of all questionnaire measures see Carhart-Harris et al. (2016).

### Stimuli and Image Acquisition

Subjects were instructed to keep their eyes closed for the duration of the resting-state and music scans. A prompt with these instructions was displayed on the screen during the scan as a reminder in case they opened their eyes at any point.

Music stimuli used in this study were edited compositions by Carlos Cipa. A different music track was played on each scan visit, and the order of the playlist was randomized across subjects. ‘Lost and Delirious’ and ‘Lie with Me’ were combined in the first track; and ‘Wide and Moving’ and ‘The Dream’ were combined in the second track. All tracks used were solo piano works, with no vocals or other instruments. The tracks were chosen in order to balance emotional potency, based on ratings by an independent sample. Ableton Live 9 software was used to boost the volume and apply audio compression in order to provide maximally-audible stimuli that could be easily heard over the background noise of the scanner. Music was played through MRI-compatible headphones (MR Confon) during the scan.

Imaging was performed on a 3T Siemens Tim Trio using a 12-channel head coil at Invicro, London, UK. Anatomical images were acquired using the ADNI-GO (Jack et al., 2008) recommended MPRAGE parameters: TE 2.98ms, 160 sagittal slices, 256?256 in-plane FOV, flip MPRAGE parameters (1mm isotropic voxels, TR=2300ms, flip angle = 9°, bandwidth = 240Hz/pixel, GRAPPA = 2).

T2*-weighted echo-planar images (EPI) for BOLD contrast were acquired for the functional scans (3 mm isotropic voxels, TR = 2000 ms, TE = 31 ms, 36 axial slices, 192 mm in-plane FOV, flip angle = 80°, bandwidth = 2298 Hz/pixel, GRAPPA = 2). Both the resting and music scans were 240 volumes or exactly eight minutes in duration. Data from other scan sequences was also collected during each session, and these have been reported previously elsewhere (Carhart-Harris et al., 2017; Mertens et al., 2020; Roseman, Demetriou, Wall, Nutt, & Carhart-Harris, 2018).

### Data Analysis

All analyses were performed using the FMRIB Software Library (FSL; v.6.03) and Analysis of Functional NeuroImages (AFNI) software. Anatomical data were processed using the fsl_anat script which involved skull-stripping and segmentation into White Matter (WM) and Cerebrospinal Fluid (CSF) masks with FMRIB’s Automated Segmentation Tool (FAST). These anatomical masks were registered to each subject’s functional space, and time-series from the pre-processed (see below) functional data were extracted to be used for later analysis.

Pre-processing of the functional data used FSL’s FEAT module and included head-motion correction, spatial smoothing with a 6mm Gaussian filter, pre-whitening and correction of auto-correlation of the time-series with FSL’s FILM algorithm, and registration to standard (MNI152) space. FEAT analysis models included the CSF and WM time-series, and six head-motion parameters (three translations, and three rotations) as regressors (Woletz et al., 2019). The purpose of these first-level analysis models was to de-noise the data by regressing out these eight time-series, and all subsequent analyses therefore used the residuals images produced by FEAT. These images were transformed into standard (MNI152) space using the parameters also derived by FEAT. The ALFF analyses were then performed on each individual subject’s pre-processed, de-noised, standard-space data using the AFNI 3dRSFC script. The data were band-pass filtered using a range of 0.01-0.1Hz in these analyses.

The resulting ALFF images were then combined in group-level analyses using FSL’s Ordinary Least Squares (OLS) mixed effects model. Results were thresholded at *Z* = 2.3, p < 0.05 (cluster-corrected for multiple comparisons). A single group mean average model (all subjects, all scans) was used to observe the overall spatial distribution of LFOs and to validate the methods. A separate group-level model was used to examine the main effect of the task (i.e. the difference between all music-listening scans, and all resting-scans). Further separate group-level models were used to compare the effect of the drug treatment/therapy within each scan type (i.e. after vs. before therapy for the rest scans, and after vs. before therapy for the music scans).

Following the recommendations of (Friston, Rotshtein, Geng, Sterzer, & Henson, 2006) and similar procedures used in (Yang et al., 2020) we defined functional Regions of Interest (ROIs) for further investigation. The first set of ROIs was defined from clusters identified in the main task effect analysis (music vs. resting scans). These were used to investigate the effect of psilocybin therapy in the clusters which showed a significant task-dependent difference. The second set of ROIs was defined from clusters resulting from the specific comparisons of before vs. after psilocybin therapy, in each functional task scan. These were used to assess relationships (using Pearson’s correlations) with appropriate psychometric measures: the 5D-ASC, the GEMS-3, and the QIDS questionnaires. All ROI data were *Z*-normalised before further analysis.

## Results

The mean (all subjects, all scans) analysis of the ALFF measure showed a regional distribution of values similar to what is typically observed in ALFF studies (e.g. Zou et al., 2008). High values were observed in ventral brain regions (likely attributable to physiological noise) but also in the cingulate cortex, medial and lateral frontal lobes, and insula (see supplementary figure S1).

Comparison of mean effects of scan type (all resting-state scans vs. all music-listening scans contrasted but pre vs post treatment collapsed into one) showed relatively higher ALFF values in the bilateral superior temporal gyrus (primary auditory cortex) for the music-listening scan, and a relatively higher response in the lingual gyrus for the resting-state scan.

These two regions were defined as ROIs and the effects of treatment (before vs. after psychedelic therapy) and scan type (rest vs. music) were examined using 2×2 ANOVA models. For the lingual gyrus region, there was only a main effect of scan type (*F*[1,18] = 38.45,*p* < 0.001), which is expected as the ROIs were selected on the basis of differential response to the two scans. In the superior temporal gyrus region, there was likewise a scan type main effect (*F*[1,18] = 32.89, *p* < 0.001) but also a significant interaction with treatment (F[1,18] = 4.90, *p* = 0.04). Post-hoc tests revealed the source of this interaction to be a significant increase in responses in this region *after* psilocybin therapy, but only for the music-listening scan (*t*[31] = 2.09, *p* = 0.045). See histograms on figure 1. This suggests a specific effect of increased LFOs in the superior temporal lobe after the therapy when listening to music. There was no effect of the therapy on the resting state scan in this superior temporal lobe region, and no effects of the therapy in the lingual gyrus region (in either scan type). Additional analyses examined potential differences between the two music tracks and the effectiveness of the counterbalancing. An additional between-subjects factor of track order was also included (i.e., subjects who received track A on visit 1, vs. subjects who received track B on visit 1). These analyses showed no main effects of track order and no significant interactions with the other factors (all *p* values > 0.1) in either of the ROIs.

**Figure 1.**
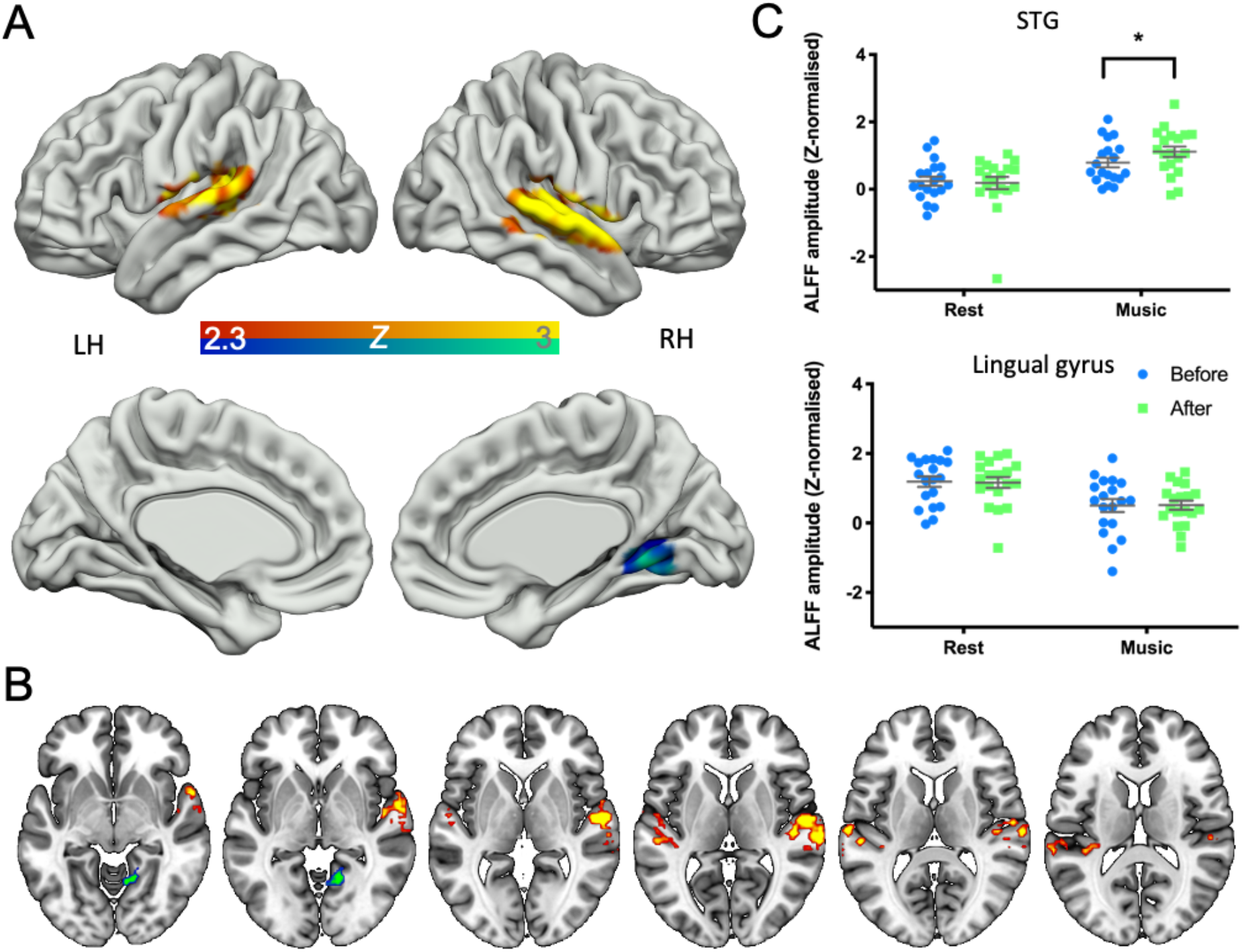
A: Comparison of the two scan types (resting-state or ‘rest’ vs. music), with pre and post treatment collapsed (panel A = 3D render, B = axial slices). The red-yellow colour-scale denotes greater responses in the music-listening (vs rest), and the blue-green colour-scale denotes greater responses in the resting scan (vs music). Panel C: ROI data from the two task-defined ROIs. Significant effect of treatment in the music/superior temporal gyrus (STG) ROI, for the music scan (*p*=0.045). Clear main effects of scan type (rest vs. music) are also evident here in both regions because these ROIs were selected on that basis.

Voxelwise comparisons of the resting-state data before vs. after treatment within each scan type (figure 2) showed an effect in the medial frontal lobe (before > after), suggesting decreased ALFF in these regions after the therapy. We also see relative increases in ALFF (after > before) in the music scan, in a lateral region covering the superior temporal areas and the supramarginal gyrus in the left hemisphere, and a similar, though somewhat smaller and more anterior region in the right hemisphere, centred on the inferior portion of the precentral gyrus. An additional small cluster in the left lingual gyrus is also present in this comparison.

**Figure 2.**
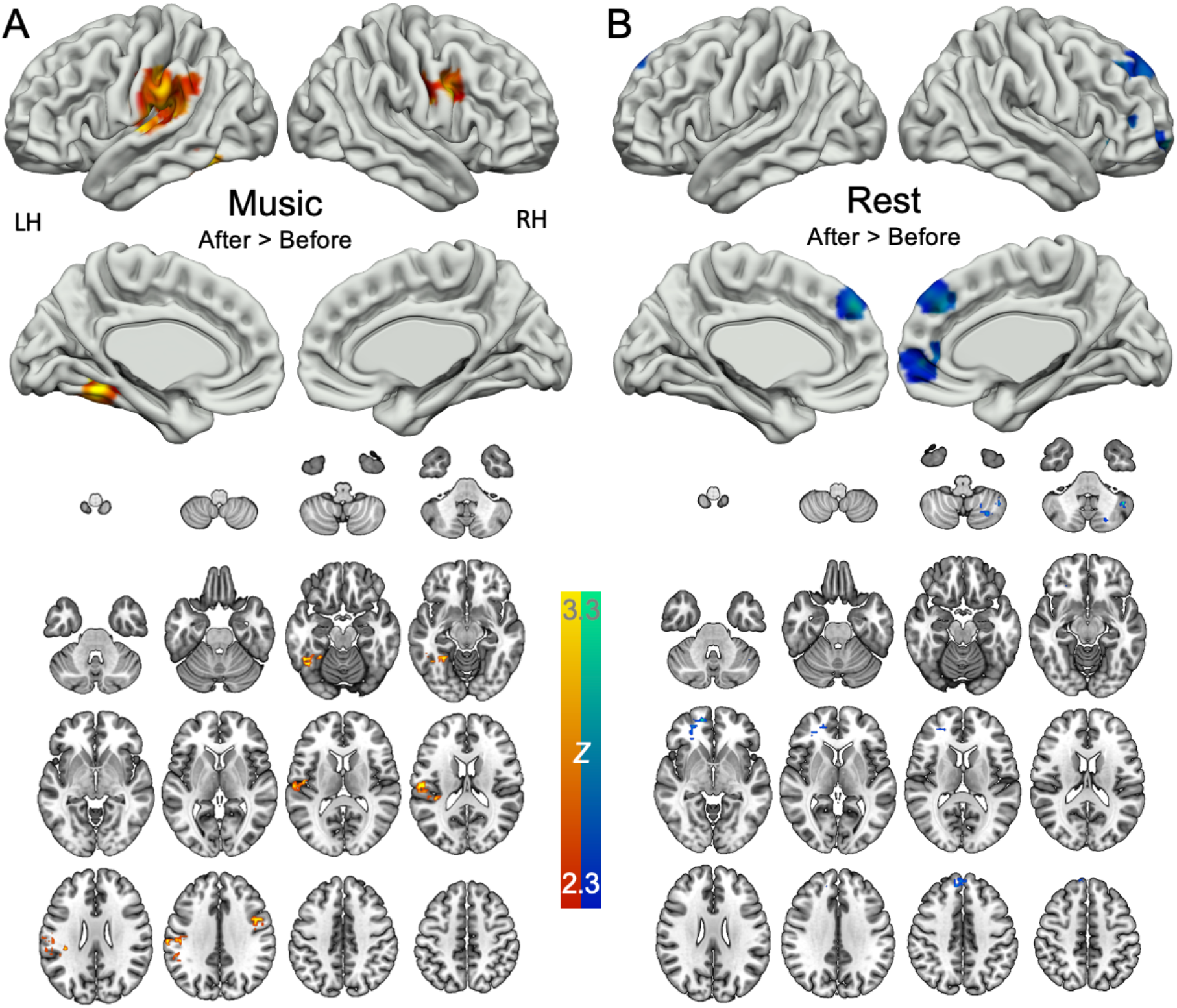
Comparison of pre- and post-therapy effects (after > before) for each scan individually (A = music; B = resting). The red-yellow colour-scale denotes increased responses post-therapy and the blue-green colour-scale denotes decreased responses post-therapy. ALFF values during the music scan were higher post-therapy with significant clusters observed in the supramarginal gyrus, extending down into the superior temporal lobe (left hemisphere), the inferior portion of the pre-central gyrus (right hemisphere), and a small cluster in the left lingual gyrus. In the resting-state scan, ALFF values were reduced post-therapy in the medial frontal lobe.

These activation clusters were also defined as ROIs and the delta (change in response between pre- and post-therapy scans: after minus before) was calculated on data from these ROIs. Performing correlations on these measures with clinical and psychometric scores showed that the change in response to music in the post-therapy scan was significantly correlated with several sub-scales of the ASC scale acquired on the second (high-dose) treatment visit (figure 3). The DED (ego-dissolution), VRS (visionary restructuralization), AUA (auditory alterations) and VIR (vigilance reduction) sub-scales showed significant relationships, while the OCEAN (oceanic boundlessness) sub-scale was non-significant. The mean of all five sub-scales was also highly correlated with the ROI data (*r* = 0.621, *p* = 0.005). The mean of all sub-scales and the result from the VIR sub-scale (*r* = 0.613, *p* = 0.005) survive family-wise correction for multiple comparisons with a corrected alpha level of *p* = 0.008. There were no other relationships between the music scan data and any other subjective questionnaire measure or clinical rating scale, and similar analyses with the rest scan data also showed no significant relationships with any of the questionnaire measures. See the supplementary material for full tables of correlation results.

**Figure 3.**
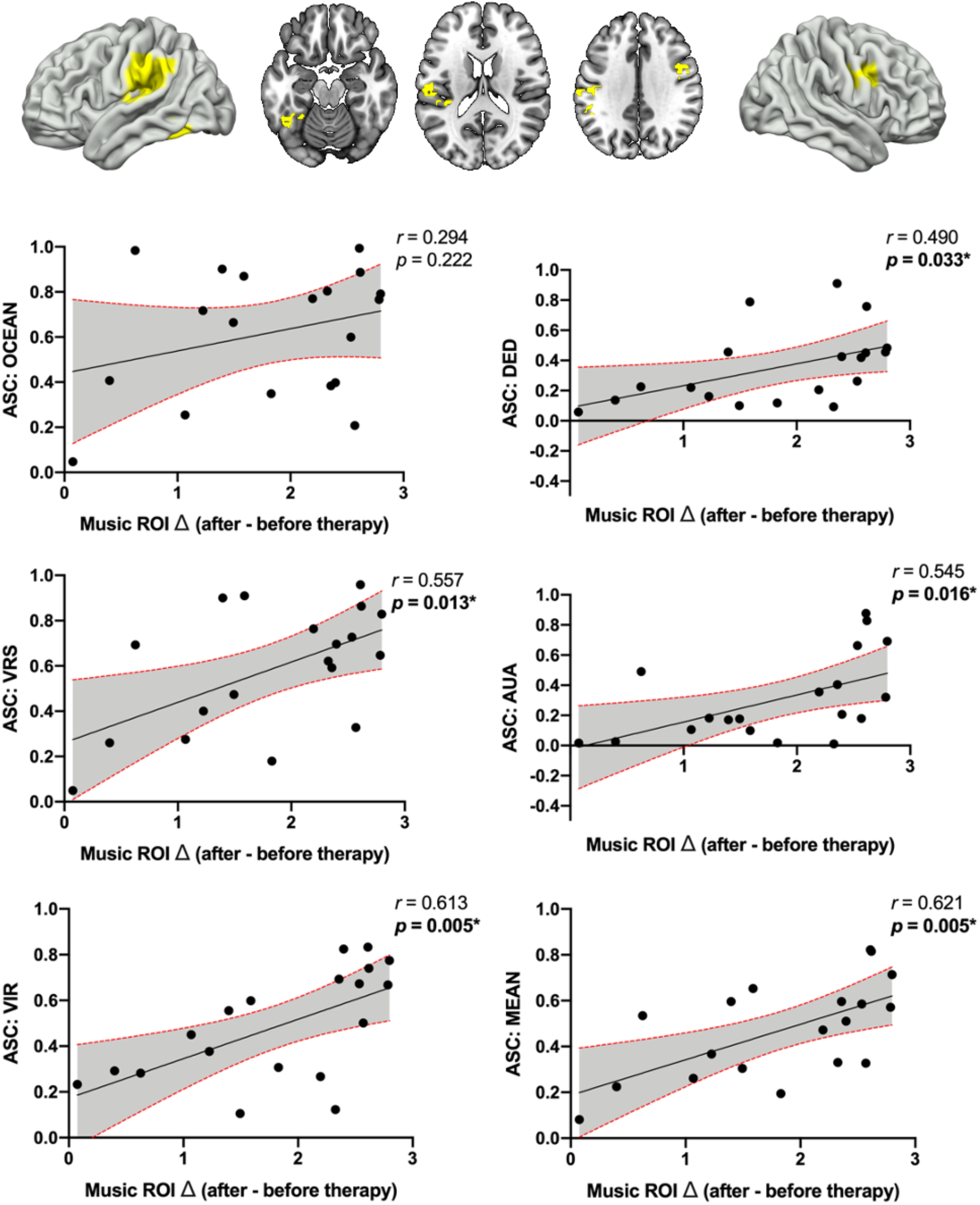
Exploratory correlational analyses between increased ALFF during music after therapy vs before (ROI: top row) and the five-dimensional altered states of consciousness sub-scales. OCEAN = “Oceanic boundlessness”; DED = “Ego-dissolution”; VRS = “Visionary restructuralization”; AUA = “Auditory alterations”; VIR = “Vigilance reduction”. MEAN = Mean of all five sub-scales (also known as the ‘global’ ASC score). Significant correlations (*p* < 0.05) are highlighted with bold text and *. The VIR and mean/global scores (bottom two panels) survive a family-wise corrected *p* value threshold of 0.008.

## Discussion

ALFF identified brain regions in the superior temporal lobe in which low-frequency fluctuations were greater during music listening compared with (no-music) resting-state conditions (figure 1) and these regions showed significantly greater increase in ALFF after psilocybin therapy versus before (figure 1 histograms; panel C). Examining the effects of the psychedelic therapy on each scan type separately revealed lower ALFF in the medial frontal lobe for the resting-state scan after therapy versus before, and increased ALFF in the superior temporal lobe regions and supramarginal gyrus for the music scan after versus before therapy. Patients therefore had higher responses in recognized music and musical emotion-processing brain regions (Koelsch et al., 2018) after the therapy. Furthermore, this increased ALFF-indexed responsiveness to music was mediated by the subjective quality of the psychedelic experience in the therapy session, with the increases in ALFF being significantly correlated with higher levels of (anxious) ego-dissolution, visionary restructuralization, auditory alterations, and vigilance reduction as well as an averaged total or ‘global’ score, all measured by the 5D-ASC questionnaire (Dittrich, 1998; Hasler, Grimberg, Benz, Huber, & Vollenweider, 2004). This suggests a potential causal effect of the drug experience in producing the effects.

The finding that music-listening (compared with rest) produces increased LFOs in the superior temporal region (Heschl’s gyrus, and the planum temporale) is unsurprising, as these regions belong to the primary auditory cortex, and are highly specialized for sound perception, including music. What is perhaps more interesting is that responses in these regions were also significantly affected by the therapy in these patients, given that most previous work on musical aesthetics and emotionality has tended to identify reward/limbic regions as being of greater importance for these features (Brown, Gao, Tisdelle, Eickhoff, & Liotti, 2011; Koelsch, 2020; Koelsch & Skouras, 2014; Menon & Levitin, 2005). However, recent work has also strongly made the case that the auditory cortex plays a role in the processing of affective auditory information, and has functional connections with limbic and paralimbic structures (Koelsch et al., 2018). A recent meta-analysis (N=47 studies) has also identified Heschl’s gyrus as being specifically involved in music-evoked emotions, as well as a range of other limbic and reward regions (Koelsch, 2020). These results therefore provide additional convergent evidence that the therapeutic effect of psilocybin is (at least, partly) mediated by the qualities of the acute psychedelic experience, including ‘emotional breakthroughs’ which are a key mediator of longer-term psychological effects, including improvements in mental health outcomes (Roseman et al., 2019). Previous work on this cohort of patients is also supportive of this interpretation, showing that changes in the functional connectivity of emotion/reward regions - such as the vmPFC - are meaningfully related to longer-term clinical effects (Carhart-Harris et al., 2017) and that brain responses to emotional face stimuli one day post-treatment show clear increases after the therapy (Roseman et al., 2018), with additional effects on brain connectivity (Mertens et al., 2020); although see Barrett, Doss, Sepeda, Pekar, & Griffiths (2020) for contrasting results after a longer period post-dose. Taken together, it is tempting to infer a greater sensitivity or responsivity to emotional stimuli in complex emotional processing systems post psilocybin therapy; consistent with findings that emotional responsiveness is enhanced post psilocybin therapy (Carhart-Harris et al., 2021).

The present study’s results also further validate the use of music as an experimental probe stimulus in studies of depression and build upon previous work in this area (e.g. Jenkins *et al*., 2018). The approach used here of a single continuous piece of music contrasted with a resting-state scan is relatively uncommon but provides a novel and rich dataset which can be interrogated with a number of different analysis approaches (Cong et al., 2014). The analysis of LFOs presented here would likely not be possible with a more conventional fMRI design (e.g. relatively short blocks of music separated by silence or non-musical sound). Use of continuous ‘naturalistic’ stimuli in fMRI is becoming more common (Breakspear & Chang, 2020; Maguire, 2012; Sonkusare, Breakspear, & Guo, 2019), with recent results showing that it may have higher test-retest reliability than standard methods (Wang et al., 2017), be more accurately predictive of behavioural phenotypes (Finn & Bandettini, 2021), and have the advantage of superior ecological validity versus e.g., short blocks of music.

The post-therapy changes seen in the resting-state scan in the medial frontal lobe are also consistent with previous work showing abnormal medial frontal lobe functioning in depression (Lemogne, Delaveau, Freton, Guionnet, & Fossati, 2012; Lemogne et al., 2009; Nejad, Fossati, & Lemogne, 2013). The medial PFC has also been implicated in the acute brain action of psychedelics (Carhart-Harris et al., 2012, 2015). Previous work using ALFF methods has identified frontal lobe abnormalities in major depression (Rosenbaum et al., 2020) and in depressed Parkinson’s disease patients (Wen, Wu, Liu, Li, & Yao, 2013). The medial frontal lobe has been implicated in the action of selective serotonin re-uptake inhibitors (SSRI) treatment (Di Simplicio, Norbury, & Harmer, 2012; Godlewska, Browning, Norbury, Cowen, & Harmer, 2016; Ma, 2015). Recent work has also shown reductions in medial-frontal connectivity with the amygdala, following psychedelic therapy (Mertens et al., 2020) as well as decreased mPFC-posterior cingulate cortex functional connectivity under psilocybin (Carhart-Harris et al., 2012) but increased mPFC-parietal lobule connectivity after psilocybin therapy for depression (Carhart-Harris et al., 2017). Taken together, these findings converge on medial PFC dysfunction in depression being a key target for psilocybin therapy.

Limitations of this study largely relate to its design as an open-label trial with a limited number of subjects and lack of placebo control; these issues will require further trials to adequately address. In the mean ALFF data (see supplementary material) the pattern of high values around the base of the brain where there are many large blood vessels, does suggest a substantial physiological (cardiac, respiratory) component in the signal, despite the de-noising procedures used. However, the experimental design used here effectively controls for these effects, with within-subjects comparisons used for both cross-session and within-session contrasts between the resting and music scans. ALFF measures have high reliability (Li et al., 2012), with physiological effects also showing high temporal stability (Küblböck et al., 2014) and are therefore effectively subtracted out by the within-subjects design. The specificity of the results of contrasts between resting and music scans (figure 1), which specifically highlight the auditory cortex, suggest that this is the case. The related measure fALFF (fractional ALFF; Zou *et al*., 2008) is somewhat less influenced by physiological effects, but also has significantly lower test-retest reliability (Zuo & Xing, 2014), making it a less suitable measure in this study, which relies on cross-session comparisons.

In summary, this study’s results suggest that naturalistic music-listening, as well as being a crucial part of the therapy itself (Barrett et al., 2018), is also a potentially useful method for investigating treatment effects in psychedelic-therapy research. Patients in this study showed an enhanced response to music-listening the day after the therapy, as indexed by increased ALFF, and this enhanced response was related to the intensity of the subjective effects felt during the acute psychedelic experience on the high-dose therapy visit. Future work may examine the effects of music with a longer gap separating the last dosing and the post treatment scan, as there are some reasons to believe that different brain changes can be seen one day post-treatment versus after a longer post-treatment period. Nevertheless, these data provide an initial indication of the effects of psychedelic therapy for depression on brain responses to music that will help to enrich our understanding of psilocybin’s therapeutic mechanisms as we go forward.

## Supplementary Material

**Figure S1.**
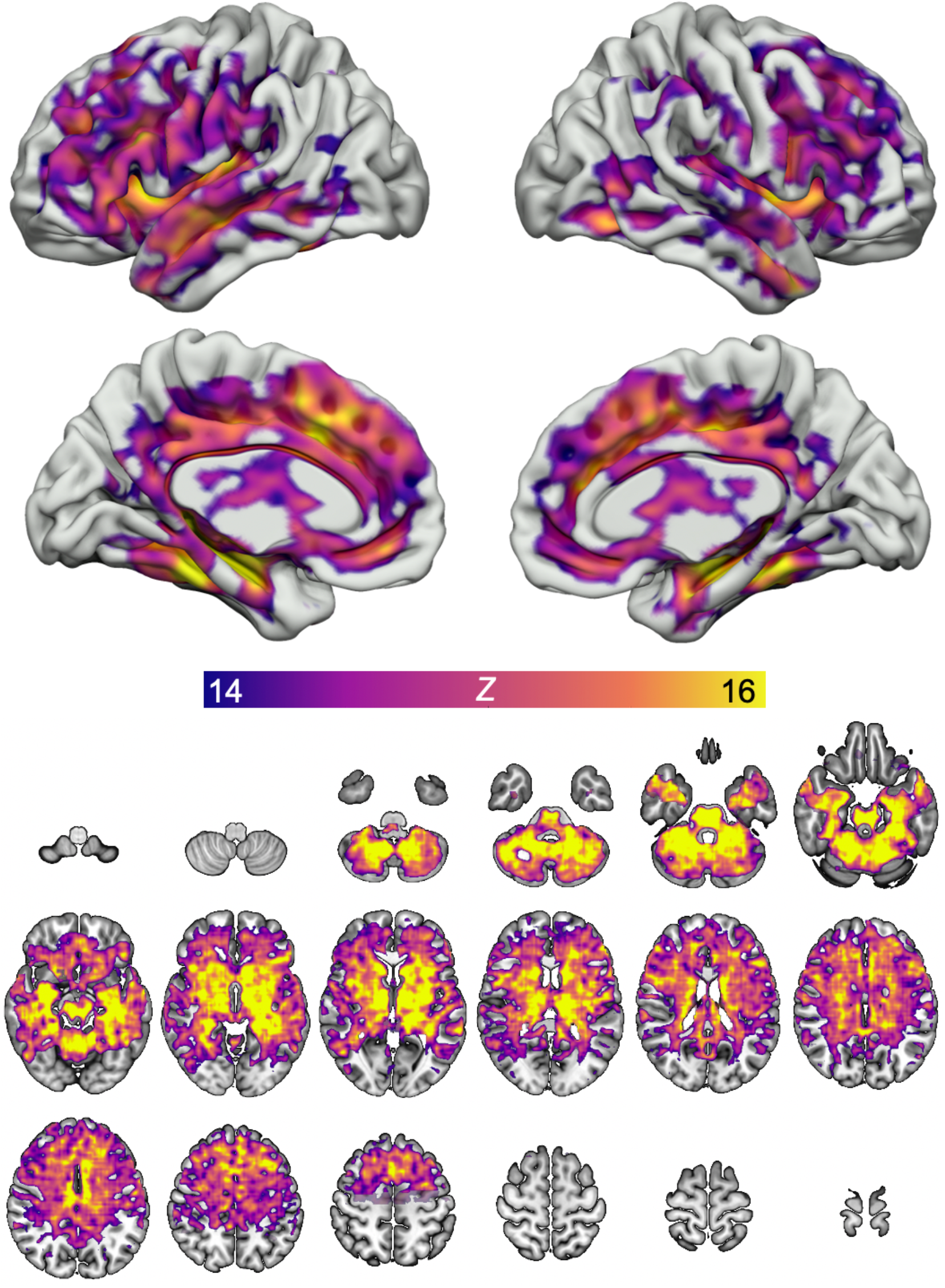
Mean ALFF values across all subjects (N=19), and all scans (four scans per subject; music-listening and resting-state scans, before and after therapy). All statistical maps are generated with cluster probability threshold of *Z*=2.3, *p*<0.05, cluster-corrected for multiple comparisons. Images are shown in neurological orientation (left hemisphere = left of the image).

**Table S1.** Correlations between the delta (after treatment – before treatment) of ALFF measures in ROIs and the GEMS-3 scale on day 2 (high-dose treatment day). All correlations were non-significant (all *p* values > 0.17).

|  | **Music $\Delta$ (after - before)** |  | **Rest $\Delta$ (after - before)** |  |
| --- | --- | --- | --- | --- |
|  | Pearson's *r* | *p*-value | Pearson's *r* | *p*-value |
| **GEMS-3, Sublimity, Day2** | 0.145 | 0.565 | 0.141 | 0.577 |
| **GEMS-3, Vitality, Day 2** | 0.336 | 0.173 | 0.141 | 0.578 |
| **GEMS-3, Unease, Day 2** | 0.079 | 0.754 | -0.069 | 0.786 |

**Table S2.** Correlations between the delta (after treatment – before treatment) of ALFF measures in ROIs and the Quick Inventory of Depression Symptoms scale at time-points ranging from baseline (start of the study) to the six-month follow-up. All correlations were non-significant (all *p* values > ~0.3).

|  | **Music $\Delta$ (after - before)** |  | **Rest $\Delta$ (after - before)** |  |
| --- | --- | --- | --- | --- |
|  | Pearson's *r* | *p*-value | Pearson's *r* | *p*-value |
| **QIDS: Baseline** | -0.209 | 0.391 | -0.005 | 0.984 |
| **QIDS: 1 Week** | -0.108 | 0.659 | -0.068 | 0.781 |
| **QIDS: 2 Weeks** | -0.06 | 0.807 | 0.058 | 0.812 |
| **QIDS: 3 Weeks** | -0.077 | 0.753 | -0.083 | 0.737 |
| **QIDS: 5 Weeks** | 0.009 | 0.97 | -0.012 | 0.962 |
| **QIDS: 3 Months** | 0.252 | 0.298 | 0.28 | 0.246 |
| **QIDS: 6 Months** | 0.009 | 0.971 | -0.026 | 0.915 |

**Table S3.** Correlations between the delta (after treatment – before treatment) of ALFF measures in ROIs and the 5-dimensional altered states of consciousness (5D-ASC) scale acquired during the second (high-dose) dosing visit. OCEAN = “Oceanic boundlessness”; DED = “Ego-dissolution”; VRS = “Visionary restructuralization”; AUA = “Auditory alterations”; VIR = “Vigilance reduction”. MEAN = Mean of all five sub-scales. Significant correlations (*p* < 0.05) are highlighted with bold text. The VIR and mean result survive a family-wise corrected *p* value threshold of 0.008.

| | Music $\Delta$ (after - before) | | Rest $\Delta$ (after - before) | |
| --- | --- | --- | --- | --- |
| | Pearson's $r$ | $p$-value | Pearson's $r$ | $p$-value |
| **5D-ASC: OCEAN** | 0.294 | 0.222 | 0.201 | 0.41 |
| **5D-ASC: DED** | 0.49 | **0.033** | 0.336 | 0.16 |
| **5D-ASC: VRS** | 0.557 | **0.013** | 0.407 | 0.084 |
| **5D-ASC: AUA** | 0.545 | **0.016** | 0.307 | 0.2 |
| **5D-ASC: VIR** | 0.613 | **0.005** | 0.406 | 0.084 |
| **5D-ASC: Mean** | 0.621 | **0.005** | 0.411 | 0.08 |

